# *C. elegans* synMuv B proteins regulate spatial and temporal chromatin compaction during development

**DOI:** 10.1101/538801

**Authors:** Meghan E. Costello, Lisa N. Petrella

## Abstract

Tissue-specific establishment of repressive chromatin through creation of compact chromatin domains during development is necessary to ensure proper gene expression and cell fate. *C. elegans* synMuv B proteins are important for the soma/germline fate decision and mutants demonstrate ectopic germline gene expression in somatic tissue, especially at high temperature. We show that *C. elegans* synMuv B proteins regulate developmental chromatin compaction and that timing of chromatin compaction is temperature sensitive in both wild-type and synMuv B mutants. Chromatin compaction in mutants is delayed into developmental time-periods when zygotic gene expression is upregulated and demonstrates an anterior-to-posterior pattern. Loss of this patterned compaction coincides with the developmental time-period of ectopic germline gene expression that leads to a developmental arrest in synMuv B mutants. Thus, chromatin organization throughout development is regulated both spatially and temporally by synMuv B proteins to establish repressive chromatin in a tissue-specific manner to ensure proper gene expression.

## Introduction

Generating proper chromatin organization during development is required for fate specification. As cells go from a totipotent one-cell zygote to a multicellular differentiated organism, chromatin transitions from being highly dynamic and non-ordered to more static, with clear euchromatic and heterochromatic domains within the nucleus (Mutlu et al., 2018; Politz et al., 2013; Yuzyuk et al., 2009). These changes in chromatin occur during a concurrent developmental transition from no/very low zygotic gene expression to zygotic, fate-specific gene expression (Levin et al., 2012; Robertson et al., 2015; Spencer et al., 2011). Chromatin compaction and accessibility are thought to have a large influence on the level of gene expression by regulating the ability of transcription factors and polymerase to have access to genes (Elgin and Reuter, 2013; Simon and Kingston, 2013). Closed or compacted chromatin domains are thought to be impenetrable by transcriptional activators and RNA polymerase and are often associated with repressed genes.

Repressive chromatin domains are formed during early embryogenesis (Gonzalez-Sandoval, et al., 2015; Motlu *et al,* 2018; Politz et al., 2013; Yuzyuk et al., 2009). There are two types of canonical repressive chromatin, constitutive heterochromatin that covers gene poor and repeat rich regions of the genome, and facultative heterochromatin that is found in gene rich regions of the genome and acts as a spatial-temporal regulator of gene repression (Ahringer and Gasser, 2018; McMurchy et al, 2017). *C. elegans* have holocentric chromosomes with gene rich, active centers and gene poor, repeat rich arms (Ahringer and Gasser, 2018; Albertson and Thomson, 1982; Gerstein et al., 2010). Constitutive heterochromatin is found along chromosome arms in *C. elegans,* and facultative heterochromatin can be found in the more euchromatic centers of chromosomes (Liu et al., 2011; McMurchy et al., 2017).

Although these classical designations of repressed chromatin have been characterized in worms, it has become increasingly clear that chromatin dynamics are complex and cannot always be classified into these two designations (Ahringer and Gasser, 2018; Liu et al., 2011). Repressive chromatin is established in active regions of the genome by tissue specific mechanisms in other organisms, but the regulators and establishment of facultative heterochromatin during crucial developmental windows for proper fate specification have yet to be elucidated (Gaertner et al., 2012; Smolko et al., 2018).

synMuv B proteins are a group of conserved transcriptional repressors important in the tissue specific repression of a large set of genes. Loss of a subset of synMuv B proteins, including members of the DREAM complex, LIN-15B, and MET-2, results in the ectopic expression of germline genes in the somatic intestine (Andersen and Horvitz, 2007; Petrella et al., 2011; Wang et al., 2005; Wu et al., 2012). Ectopic germline gene expression in the intestine leads to High Temperature L1 larval Arrest (HTA) in DREAM complex and *lin-15B* mutants (Petrella et al., 2011). The DREAM complex and LIN-15B bind to the promoter region of target genes throughout the genome where they are thought to repress their expression. The highly conserved DREAM complex has eight known subunits including LIN-35 and LIN-54 (Fay and Yochem, 2007; Harrison et al., 2006). LIN-35, the single worm homolog of the mammalian pocket protein retinoblastoma, mediates the interaction of the subcomplex portions of the DREAM complex and in it its absence the DREAM complex has highly reduced binding and repression capabilities (Goetsch et al., 2017). LIN-54 acts as one of the DNA binding subunits of the DREAM complex and in its absence the DREAM complex is lost from 80-90% of target loci (Tabuchi et al., 2011). LIN-15B has not been shown to be part of the DREAM complex, but demonstrates the same phenotypes as seen in DREAM complex mutants, including ectopic germline gene expression and HTA (Petrella et al., 2011). MET-2 catalyzes mono- and di-methylation of histone H3 lysine 9 (H3K9me1 and H3K9me2), a histone modification associated with repressive chromatin (Andersen and Horvitz, 2007; Towbin et al., 2012). *met-2* mutants lose ~80% of H3K9me2 (Andersen and Horvitz, 2007; Towbin et al., 2012). Although loss of the DREAM complex, LIN-15B, and H3K9me2 catalyzed by MET-2 all lead to ectopic expression of germline genes, the role and order of each of these proteins during development remains unknown.

The phenotypes demonstrated by synMuv B mutants are temperature sensitive. Larval arrest only happens at high temperature and ectopic germline gene expression is more extensive at high temperatures. Chromatin has been shown to be affected by changes in temperature in other organisms. For example, early studies in *Drosophila* revealed that high temperature causes incomplete polytene heterochromatin formation (Hartmann-Goldstein, 1967). Additionally, work in plants has shown that flowering time is linked to changes in chromatin that occur in response to increased temperature (Zografos and Sung, 2012). Therefore, the temperature sensitivity of synMuv B mutants may be associated with temperature sensitivity of chromatin. We hypothesized that synMuv B proteins regulate gene expression at the chromatin level throughout development and are particularly necessary for the proper chromatin regulation and buffering of gene expression during temperature stress.

Here we investigate the loss of synMuv B proteins on chromatin compaction and accessibility during development and during times of temperature stress. The temperature sensitivity of chromatin in *C. elegans,* and its regulating proteins, has never been examined in a developmental context before this study. We report that the timing of chromatin compaction during embryogenesis is delayed at high temperatures, even in wild-type embryos. We demonstrate that synMuv B mutants have an increased amount of open chromatin, both in general chromatin assays and at specific germline gene loci, especially at high temperature. We find the delay in chromatin compaction in synMuv B mutants occurs in an anterior posterior pattern. This patterned developmental delay occurs during the temperature sensitive periods for the high temperature arrest phenotype and ectopic germline gene expression. Our data support a mechanism that synMuv B proteins are necessary to regulate the proper spatial and temporal formation of repressive chromatin throughout development to promote fate-specific gene expression programs.

## Results

### synMuv B proteins regulate developmental chromatin compaction

To determine if chromatin compaction is compromised in synMuv B mutants, we utilized the Nuclear Spot Assay to visualize chromatin compaction during development (Yuzyuk et al., 2009). The Nuclear Spot Assay uses an extrachromosomal array containing numerous *lacO* sites that are bound by a ubiquitously expressed GFP-LacI protein.

This assay has been previously utilized in *C. elegans* to visualize chromatin compaction throughout development as differentiation is achieved (Gonzalez-Sandoval et al, 2015; Yuzyuk et al, 2009). We predicted that if synMuv B proteins are important for establishing chromatin compaction during development, we would see more open chromatin in mutants at a later stage of development as compared to wild-type worms.

We visualized chromatin compaction in intestinal cells in wild-type, *lin-15B, lin-35, lin-54,* and *met-2* mutants at three embryonic stages and in L1 larvae at 20°C and 26°C. These specific embryonic stages were chosen based on the state of zygotic gene expression and ease of staging. First, 8E embryos have eight intestinal cells (~100 cell embryo) and represent an early embryonic stage where the zygotic gene expression program has yet to be fully activated. Second, 16E embryos have sixteen intestinal cells (~200 cell embryo) and represent an early-to-mid embryonic stage when zygotic gene expression is starting to be upregulated. Finally, comma stage embryos have twenty intestinal cells and represent a mid-to-late embryonic stage when zygotic gene expression is fully underway (Fig 1A). We scored intestinal cells as having either open or closed arrays to determine differences between strains and temperatures (Fig. 1B) (Yuzyuk et al., 2009). In wild-type embryos we observed that at 20°C by the 8E stage, chromatin was already in a primarily compact state in intestinal cells and stayed compacted throughout the rest of development (Fig. 1B). However, in wild-type at 26°C at the 8E and 16E stages, we saw that there were significantly more intestinal nuclei that displayed open arrays, but that by comma stage and later the arrays were closed (Fig. 1B). This suggests that temperature causes a delay in developmental chromatin compaction even in a wild-type state.

**Figure 1:**
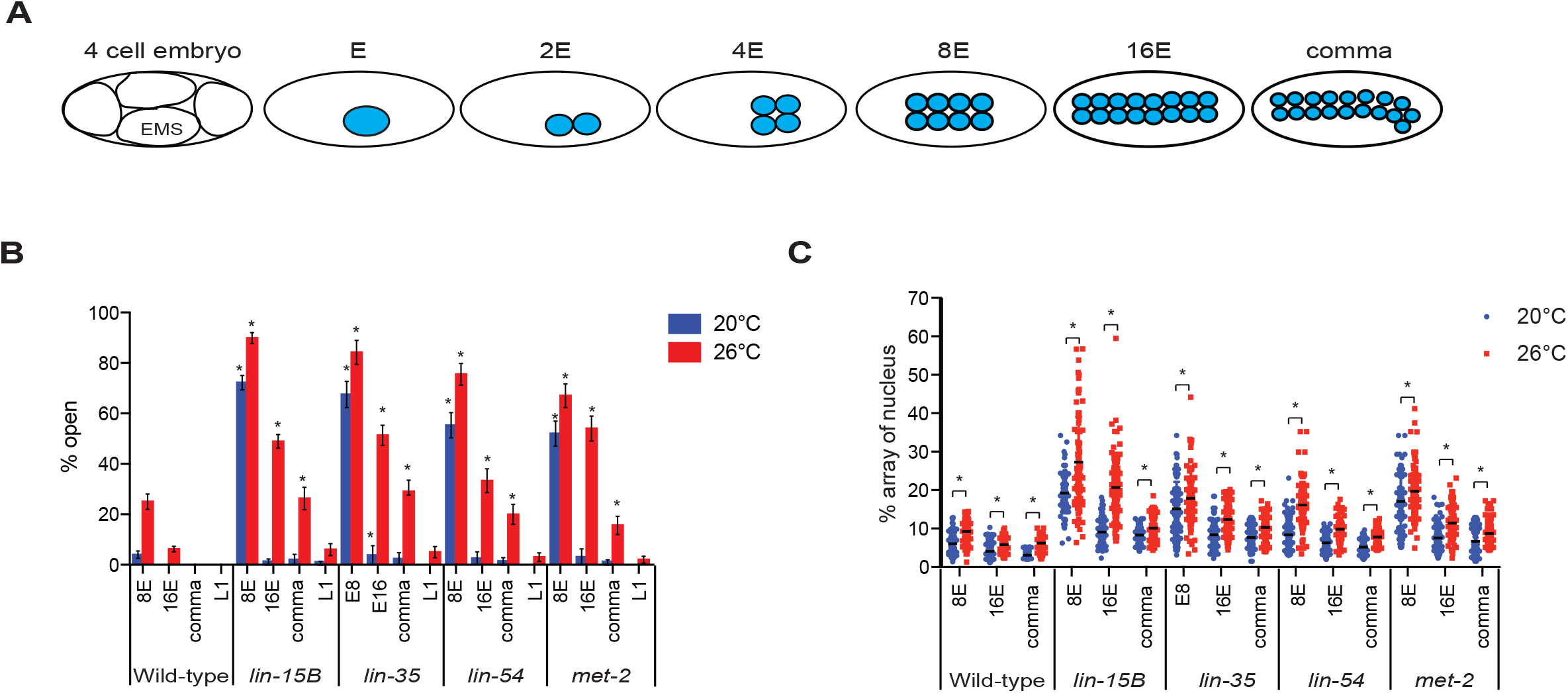
Loss of synMuv B proteins causes a delay in chromatin compaction in the intestinal lineage during development. (A) A cartoon representing the number of intestinal cells and the approximate number of total cells in the stages scored throughout this study. (B) Large extrachromosomal arrays containing numerous *lacO* sites represents a pseudo chromosome that when bound by LacI::GFP is used to visualize chromatin morphology. Intestinal nuclei were scored as open or closed based on array morphology (Yuzyuk et al., 2009). L4 hermaphrodites were placed at 20°C or 26°C and mixed stage F1 embryos were collected and percentage of intestinal nuclei with open arrays at the 8E, 16E, comma, and L1 stage at 20°C and 26°C were scored. 2-way ANOVA was used to determine significance: Asterisks represent significant difference between the wild-type population at that stage and temperature (p<0.01). Error bars indicate standard error of the proportion. n>100 cells (C) 3-D volumetric chromatin compaction analysis was performed using the isosurface function in Metamorph. The volume of the nucleus and the array were measured and used to calculate the percentage of the nucleus that the array encompassed. The percentage of the nucleus contained by the array was plotted at 8E, 16E, and comma at 20°C and 26°C. Each dot represents a single intestinal cell and the line represents the median. Asterisk represents a significant difference between 20°C and 26°C (2-way Anova p<0.01).

In all four synMuv B mutants at 20°C at the 8E stage, we saw that there were significantly more intestinal nuclei with open arrays compared to wild-type embryos at 20°C (Fig. 1B). However, for all synMuv B mutants at 20°C by the 16E stage, there were very few intestinal cells with open arrays, and only in *lin-35* mutants was the number of cells with open arrays significantly higher than wild-type at 20°C (Fig. 1B). In mutants at 20°C a small number of intestinal cells did have open arrays into comma stage but it is not significantly more than was seen in wild-type (Fig. 1B). At 26°C, mutants show a more drastic delay in chromatin compaction displaying open arrays through all stages. In all four synMuv B mutants at 26°C at all stages we saw that there were significantly more intestinal nuclei with open arrays compared to wild-type embryos, with the exception of *lin-54* and *met-2* mutants at the L1 stage (Fig. 1B). Furthermore, we saw that within an individual mutant background at 26°C compared to 20°C, there were significantly more intestinal nuclei with open arrays at each stage of development, with the exception of *met-2* mutants at the L1 stage (Fig. 1B). Therefore, synMuv B mutant intestinal cells showed a developmental delay in compaction at 26°C compared to 20°C, similar to wild-type intestinal cells. Additionally, mutants showed a higher percentage of cells with open arrays at each stage; and therefore, in synMuv B mutants at high temperature, compaction is delayed into developmental stages where zygotic gene expression has started.

To confirm our scoring method, we performed 3-D volumetric analysis of the intestinal nucleus measuring the volume of an array compared to the volume of the nucleus. Open arrays take up more volumetric space; and therefore, have a larger array volume to nuclear volume ratio. 3-D volumetric data showed similar trends to those described above; the array volume as a percent of nuclear volume was larger for mutant than wild-type, and at 26°C than at 20°C (Fig. 1C). In contrast to the open/closed scoring in wild-type at 26°C compared to 20°C, even at stages where we scored most or all of the arrays as closed, we saw a small but significant increase in the nuclear percentage of the array (Fig. 1C). This indicated arrays take up a variable amount of nuclear volume even when they appear compacted. In mutants, there was a striking increase not only in the percent of nuclear volume taken up by the array compared to wild-type, but also an increase in the variance of the array volume within the same genotype especially at 26°C. In general as development proceeded, mutant cells show a decreased percentage of array, decreased variance of array volume within a genotype, and a significant difference between temperatures (Fig. 1C). Overall, the 3-D volumetric analysis supports our open/closed analysis data as we see a temperature sensitive chromatin compaction phenotype throughout development, predominantly in synMuv B mutants.

### Individual embryos contain both nuclei with open chromatin and closed chromatin

While scoring array compaction we observed that population wide, the number of intestinal cells with open arrays decreases gradually through development in mutants. For example, in pooled *lin-15B* mutant intestinal cells at 26°C at the 16E stage about 50% of the cells contain open arrays (Fig. 1B). We wanted to determine if, at a particular stage of development, the partial compaction seen in the population of intestinal cells was due primarily to variance in compaction of individual intestinal cells within an embryo (Fig. 2A: Model 1) or variance in compaction of intestinal cells between different embryos (Fig. 2B: Model 2). If model 1 is supported, we would expect to see a population of embryos with a mosaic of open and closed cells within each embryo. If model 2 is supported, we would expect to see a population of embryos with a binary distribution of embryos containing all intestinal cells open or all intestinal cells closed. We found that in a population of embryos, each embryo contained a mix of intestinal cells with both open and closed arrays that was close to the population mean (Fig. 2C). For example, in *lin-15B* mutants at 26°C at 8E, when about 90% of intestinal cells were scored as open (Fig 1B), embryos have a range of 6/8 to 8/8 cells with open arrays. While at 16E, when about 50% of intestinal cells were scored as open (Fig 1B), *lin-15B* mutant embryos have between 6/16 and 10/16 cells with open arrays. By comma stage, *lin-15B* mutants show decreased numbers of open cells with a range of 2/20 and 6/20 cells scored with open arrays (Fig 2C). We did not find in any genotype or temperature with a bimodal distribution where the embryos within a population fell into a mix of 100% closed and 100% open arrays. This data supports model 1. As development proceeds and chromatin compaction is achieved, the number of open cells within an embryo decreases. The pattern of compaction within an embryo is similar between mutant genotypes at the same stage and temperature (Fig 2C). Mutants do display differences in mean number of open cells, but all embryos have nuclei with both open and closed arrays.

**Figure 2:**
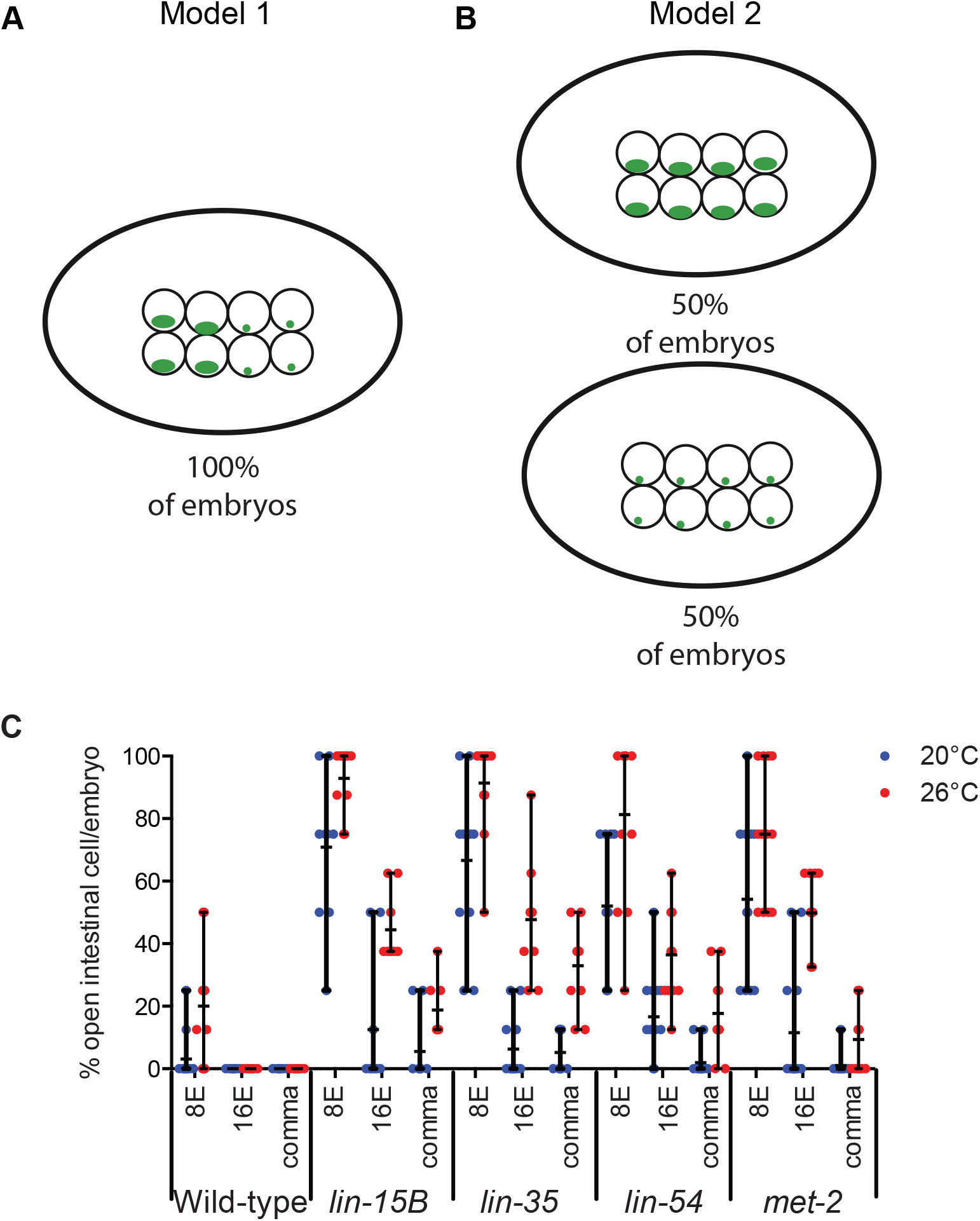
The number of intestinal cells with open chromatin varies within a single nucleus. Models that explain (A) a variance in compaction of intestinal cells within an embryo or (B) a variance in compaction of intestinal cells between different embryos. (C) The percentage of intestinal cells with an array in an open configuration per embryo was determined for 8E, 16E and comma stage embryos at 20°C and 26°C. Each dot represents an embryo and the line represents the mean percentage of open cells per embryo.

### synMuv B proteins regulate chromatin compaction spatially in an anterior-to-posterior direction

While scoring the number of open cells within an embryo, we observed an anterior-to-posterior pattern of chromatin compaction. The intestinal cell lineage descends from a single cell, E. The first cell division is an anterior-posterior division in which E becomes 2E. These two intestinal cells divide in a left-right division (2E to 4E) to establish the bilateral symmetry of the intestine. The resulting cells are designated as sister cells with a set consisting of a left and right pair. The third division when 4E gives rise to 8E is an anterior-posterior division although the cells are slightly displaced in the dorsal-ventral plane (Fig 3A, sister cells labeled as a′, c′, e′, f′). The 4^th^ division is also an anterior-posterior division that gives rise to 16E with some cells more dorsal than others (Fig 3B, sister cells labeled as a, b, c, d, e, f, g, h). The sixteen cells subsequently polarize and change shape before there is a final round of cell division in which only the most anterior pair of sister cells (a) divide in a dorsal-ventral direction, and the most posterior pair of sister cells (h) divide in an anterior-posterior direction leading to the final arrangement of cells within the nucleus (Fig 3C, sister cells labeled a, a’’, b, c, d, e, f, g, h, h’’). We found that there was a clear pattern in the anterior-posterior location of cells with open arrays in an embryo. At 8E when mutants have varying numbers of cells open at 20°C and 26°C, we found that the anterior pair of sister cells maintained open arrays more often in all four genotypes, while the most posterior cells most often had closed arrays (Fig 3). For example, at 20°C, when *lin-15B* mutants have a range of 2/8 to 8/8 cells open, the a’ sister cells were open 100% of the time. The following pair of sister cells, f’, had the next highest percentage of open cells (90%). The more posterior sister cells, c’, have a decreasing number of open cells (60%), and the two most posterior sister cells, h’, have the lowest percentage of open cells (30%) (Fig. 3A). This pattern of open arrays being more often in the anterior and less often in the posterior was observed in all four mutants, at 8E, 16E, and comma stages, at both 20°C and 26°C (Fig 3). Overall if an embryo had any intestinal cells with an open array, they were always the cells found at the anterior section of the intestine (Fig 3). This suggests that as development proceeds and chromatin compaction is achieved, cells close in a posterior to anterior direction. This pattern seems to be independent of proliferative state, as there is a final cell division at both anterior and posterior positions to give rise to 20E (Figure 3C a and a’’, h and h’’)

**Figure 3:**
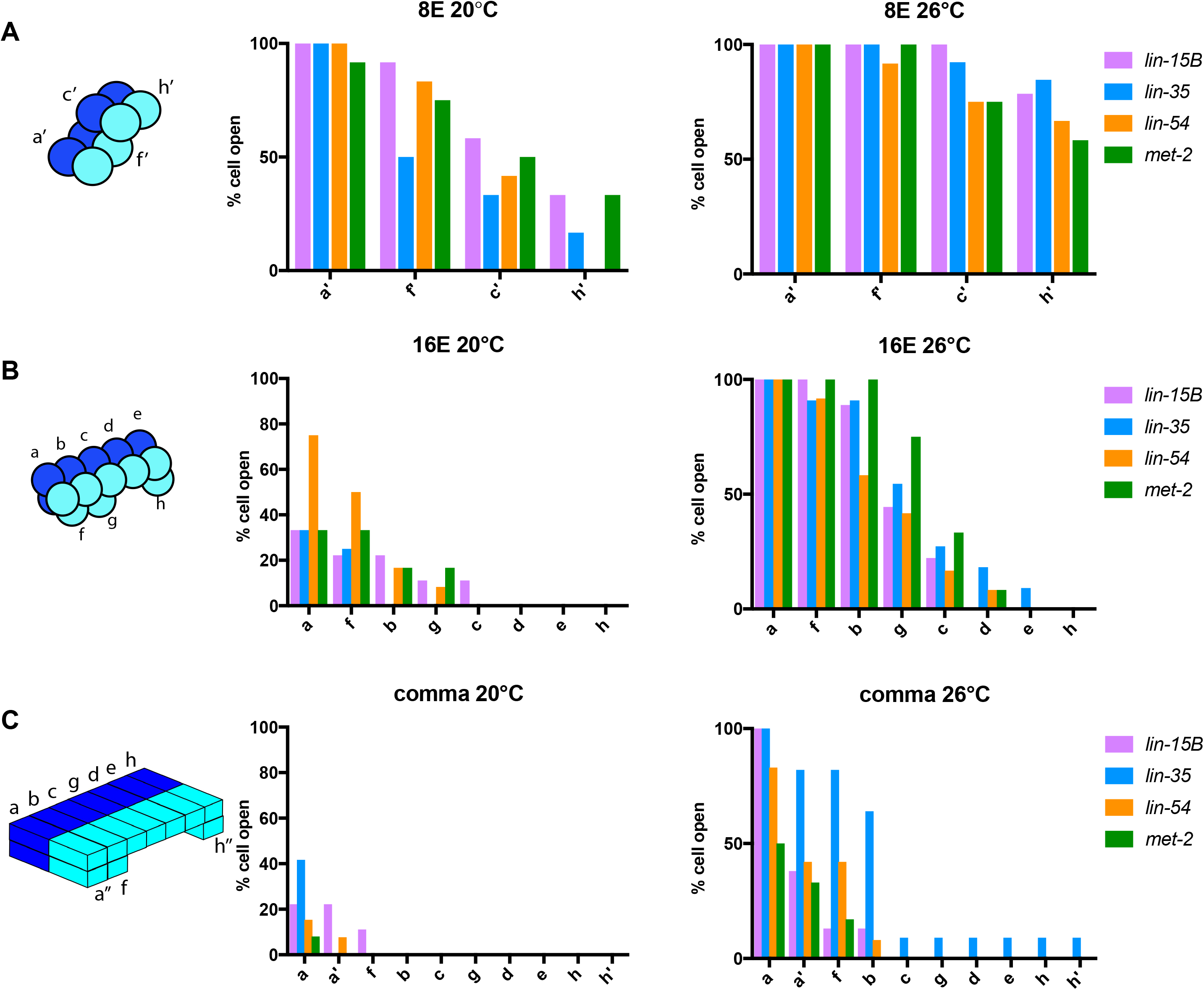
In synMuv B mutants anterior intestinal cells are more delayed in compaction than posterior intestinal cells. The position of open cells within a single embryo was monitored throughout development at (A) 8E, (B) 16E, and (C) 20E at both 20°C and 26°C. The anterior-to-posterior location of a cell was noted and that cell was recorded as either open or closed. The percent open of each pair of sister cells was plotted to in an anterior-to-posterior directionality.

### synMuv B proteins regulate compaction of endogenous germline gene loci

To determine if chromatin compaction is also altered at endogenous loci in synMuv B mutants, we performed fluorescent *in-situ* hybridization (FISH). Pairs of red and green probes ~100kb apart were created to flank endogenous loci to determine the chromosomal level of compaction at those loci (Yuzyuk et al., 2009). All loci assessed were located in central regions of autosomes that are generally euchromatic to control for autosomal location. As controls we used probes that flank the *pha-4* gene that should be open and expressed in the intestine, and probes that flank the *myo-3* gene that should be closed and not expressed in the intestine (Ardizzi and Epstein, 1987; Horner et al., 1998,). We investigated three synMuv B regulated loci *ekl-1, coh-3,* and C05C10.7. All three synMuv B target genes are germline genes that are ectopically upregulated in synMuv B mutants, and whose expression is downregulated upon rescue of high temperature larval arrest (Petrella et al., 2011). Intestinal cells were imaged in 3D stacks at the 8E, 16E, and comma stage, and we calculated the level of chromatin compaction of the loci based on the 3-D distance between the centroid of each probe (Yuzyuk et al., 2009).

To understand how compaction of endogenous loci changes with respect to expression status in the intestine and temperature, we looked across developmental time and at different temperatures in mutants and wild-type. We found that, as predicted, the nonexpressed *myo-3* control locus was more compact than the expressed *pha-4* control locus in all genotypes at both temperatures in all embryonic stages (Fig. 4A and B, Fig. S1). The *myo-3* locus demonstrated compaction reminiscent to compaction of extra-chromosomal arrays, which are also not expressed (Fig. 4A and B, Fig. S1). In wild-type at 20°C *myo-3* is completely compacted by the 8E stage (Fig. 4A and B, Fig. S1A). However, in wild-type at 26°C, there was a significant difference in the level of compaction of the *myo-3* locus between the 8E and 16E stages, representing a delay in compaction timing compared to 20°C (Fig. 4A and B, Fig. S1A). Thus, although there may be slight delays in chromatin compaction of an unexpressed locus at elevated temperatures or in mutants, cellular mechanisms can overcome these delays by late embryogenesis.

**Figure 4:**
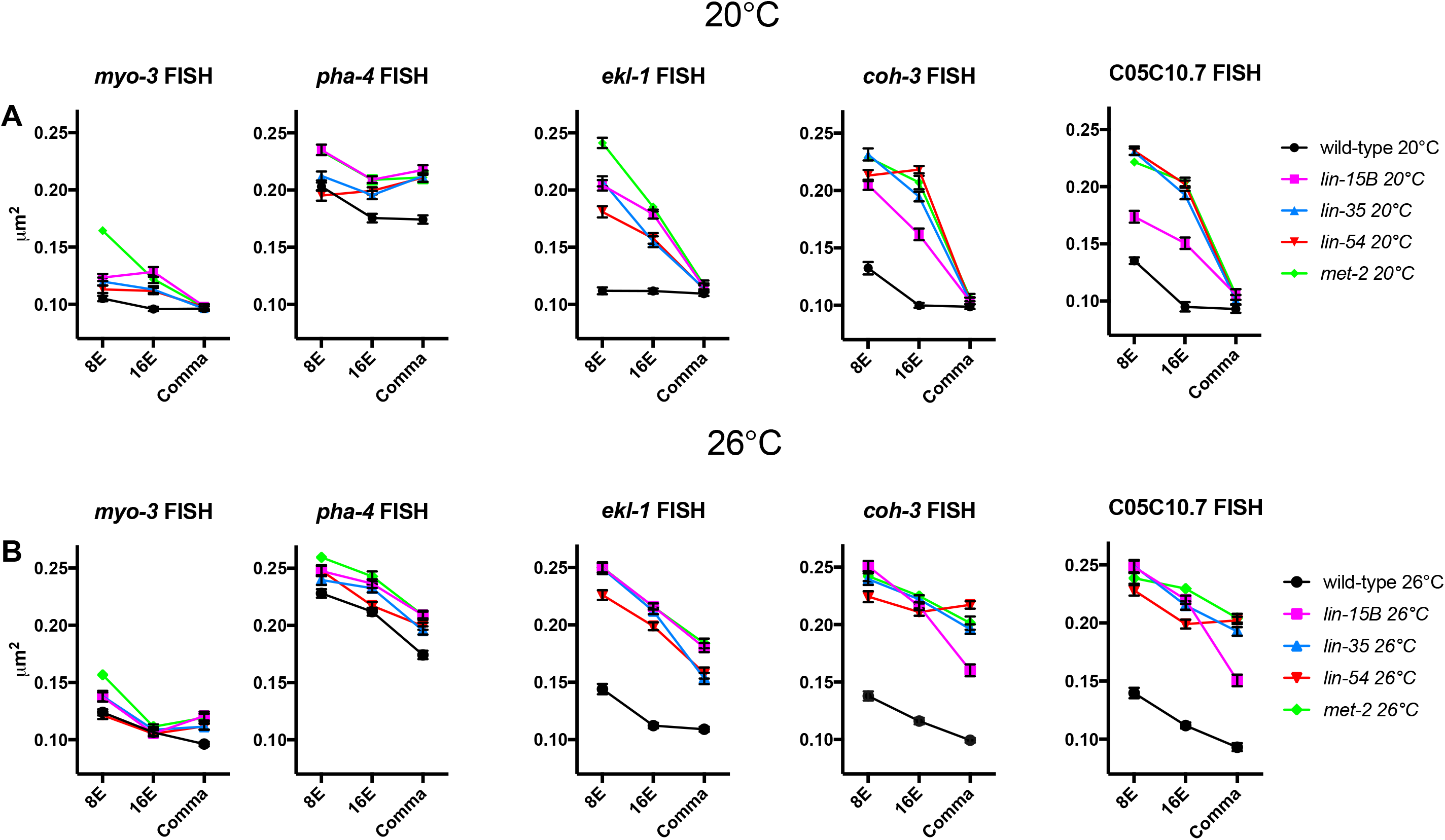
Fluorescent *in-situ* hybridization of synMuv B regulated loci reveals more open chromatin in synMuv B mutants. 8E, 16E, and comma stage wild-type, *lin-15B, lin-35, lin-54,* and *met-2* embryos at (A) 20°C and (B) 26°C were labeled with 568-5-dUTP (red) and Alexa Fluor 488 (green) probes 50kb upstream and 50kb downstream of *myo-3, pha-4, ekl-1, coh-3* and C05C10.7. Plots represent the distribution of 3-D-distances between centroids (μm^2^). See also Figure S1

We next analyzed the three synMuv B target genes *(ekl-1, coh-3, C05C10.7),* which have higher expression in mutants than wild-type and higher expression at 26°C within mutants. In wild-type, all three of these loci showed similar patterns to the *myo-3* locus with all three loci reaching maximal compaction between the 8E or 16E stage at both temperatures (Fig. 4A and B, Fig. S1A). Thus, in wild-type at both 20°C and 26°C the synMuv B target loci showed a pattern of compaction like a non-expressed locus. However, in synMuv B mutants the three targets genes showed two different patterns of developmental compaction depending on the temperature being assayed. In synMuv B mutants at 20°C at the 8E stage the *ekl-1, coh-3,* and C05C10.7 loci were all less compacted than the same locus in wild-type (Fig. 4A). However, in synMuv B mutants at 20°C by comma stage all three loci were equally compacted compared to the same locus in wild-type (Fig. 4A). Thus, at 20°C the three synMuv B regulated loci showed a delay in compaction but by late embryo genesis were as compact as in wild-type. On the other hand, in synMuv B mutants at 26°C the three synMuv B target loci were more open than the same locus in wild-type at all developmental stages (Fig. 4B). In *lin-35, lin-54,* and *met-2* mutants, both the *coh-3* and C05C10.7 were as open as the *pha-4* locus in the same genotype (Fig. 4B, Fig S1C, D, E). Thus, synMuv B target loci demonstrate a level of chromatin compaction similar to highly expressed genes late into development, but only at 26°C. The presence of open chromatin around synMuv B target genes during later developmental time periods at high temperature may leave these genes vulnerable to the ectopic misexpression that underlies the synMuv B mutant.

### 16E through comma stage is the critical developmental time-period for larval high temperature arrest in synMuv B mutants

synMuv B mutants arrest at the L1 larval stage, which is at least in part due to ectopic expression of germline genes (Petrella et al., 2011). To determine if the developmental time-period important for the HTA phenotype is consistent with embryonic stages in which we see prolonged open chromatin, we performed temperature shift experiments on *lin-15B, lin-35, lin-54,* and *met-2* mutants. Previous studies determined that the temperature-sensitive period for HTA is in a broad developmental range during embryogenesis and early larval development (Petrella et al., 2011). Since we are seeing distinct differences in chromatin compaction at different stages of embryonic development, we set out to determine a more precise time-period during embryogenesis crucial for HTA.

To perform HTA experiments, 4-cell embryos expressing a fluorescent intestinal cell marker were dissected from gravid adults and downshifted or upshifted at 4E, 8E, 16E, comma, pretzel and L1 developmental stages. Pretzel stage was included as an easily distinguishable later developmental stage to tease apart differences seen in worms shifted at comma and L1. Wild-type embryos were also dissected and shifted at all stages and never arrested (data not shown). All mutant embryos upshifted to 26°C from 20°C at 4E or 8E arrested at the L1 stage (Fig. 5A, B, C, D). When upshifted at 16E all mutant genotypes except *lin-15B* showed a slightly fewer arrested animals than at earlier stages (Fig. 5A, B, C, D). A smaller number of embryos upshifted at comma and pretzel stages arrested (Fig. 5A, B, C, D). Finally, embryos upshifted at the larval L1 stage did not arrest (Fig. 5A, B, C, D). Together, embryos shifted early in development arrested whereas embryos shifted in mid-to-late embryogenesis sometimes arrested, suggesting the critical time period for HTA to be after 16E. To complement this experiment, we also performed downshift experiments to hone in on the developmental window crucial for HTA. Embryos downshifted from 26°C to 20°C at 4E did not arrest and about half of embryos downshifted at 8E arrested (Fig. 5A, B, C, D). Strikingly most embryos downshifted at 16E and comma arrested (Fig. 5A, B, C, D). And downshifting at pretzel and L1 stages caused 100% arrested at the larval L1 stage (Fig. 5A, B, C, D). These data combined suggest 16E through comma is the most crucial temperature sensitive time-period for HTA. However, upshifting after or downshifting before this time-period can still cause some animals to undergo HTA, suggesting a buffering in the system that may represent stochastic chromatin compaction, varying amounts of ectopic gene expression, and a leaky arrest phenotype.

**Figure 5:**
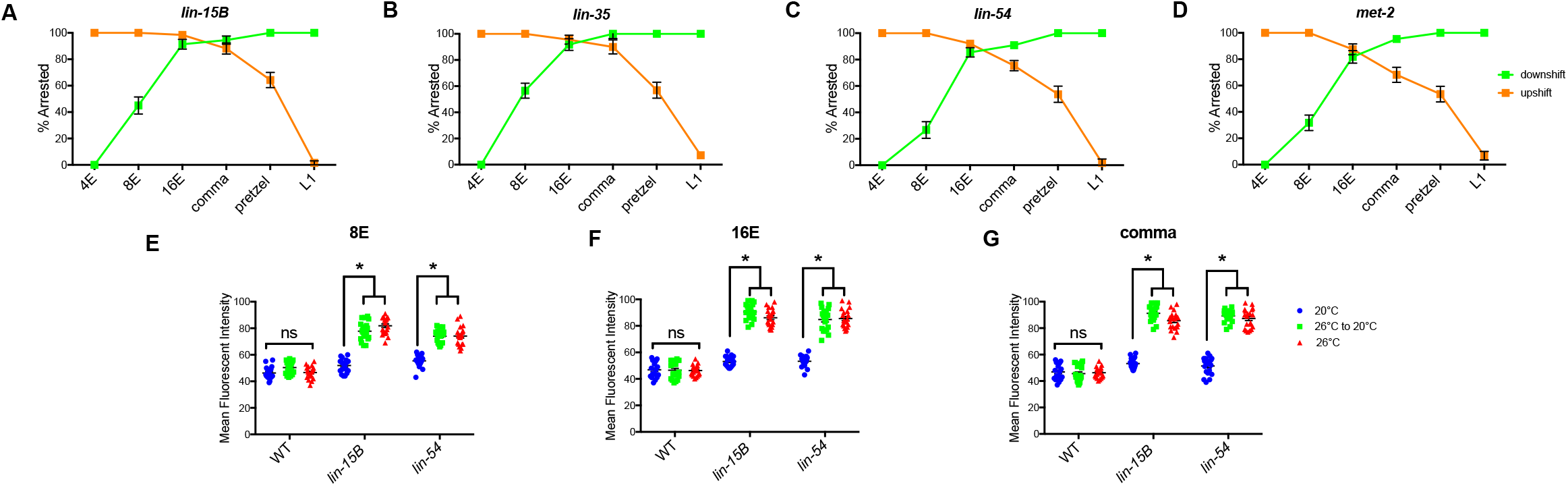
16E through comma stage of embryogenesis is the temperature sensitive time period crucial for larval HTA. (A-D) *lin-15B* (A), *lin-35* (B), *lin-54* (C), and *met-2* (D) L4 P0 hermaphrodites were placed at 20°C for upshift or 26°C for downshift experiments. F1 two cell embryos were dissected from P0s, and upshifted (red) or downshifted (blue) at 4E, 8E, 16E, comma, pretzel or L1 and scored for L1 larval arrest (n=60-78/point). (E-G) Mean Fluorescent Intensity of PGL-1 expression was determined for L1 wild-type, *lin-15B,* and *lin-54* embryos that were downshifted at 8E (E), 16E (F), comma (G) (n=20 L1/stage and temperature). 2-way ANOVA used to determine significance between samples (p<0.01). See also Figure S2

Downshift from 26°C to 20°C at pretzel and L1 stages resulted in 100% larval arrest (Fig. 5A, B, C, D), suggesting that the ectopic gene expression programs leading to HTA are activated and irreversible. Although it was previously reported that HTA was irreversible upon downshifting, whether germline genes continued to be ectopically expressed after downshift was not assessed (Petrella et al., 2011). We set out to determine if the germline gene *pgl-1* continues to be ectopically expressed after downshift by using strains with the endogenous *pgl-1* locus tagged with GFP (Andralojc, et. al., 2017). *pgl-1* is a synMuv B regulated germline gene that has been used to show ectopic germline gene expression in synMuv B mutants (Petrella et al., 2011, Wang et al., 2005, Wu et al., 2012). *lin-15B* and *lin-54* mutant embryos were downshifted at 8E, 16E, and comma stage and imaged at the L1 stage to determine the level of ectopic PGL-1 expression. We found that downshifted embryos resembled embryos that were not downshifted and kept at 26°C at all stages (Fig. 5E, F, G). There was no difference in the maximum fluorescent intensity when comparing downshifted embryos to those that remained at 26°C (Fig. 5E, F, G). There was a significant difference when comparing shifted embryos and those kept at 26°C to mutant embryos raised at 20°C (Fig. 5E, F, G). synMuv B mutants also show greater ectopic PGL-1 expression when compared to wild-type at the same stage and temperature, including 20°C (Fig. 5E, F, G). This suggests that the ectopic gene expression program that leads to HTA is activated in early embryogenesis and remains ectopically activated even upon relief of the stress.

## Discussion

Chromatin compaction during development is necessary to achieve proper gene expression and differentiation. Previous studies have identified that H3K9 methylation is essential for chromatin compaction and organization (Ahringer and Gasser, 2017; Towbin et al., 2012). Although methylation of H3K9 is necessary, the pathways and proteins needed to establish areas of compaction in a tissue specific way remain unclear. In this study we show that synMuv B proteins are necessary for timely chromatin compaction during development. Loss of any of these proteins; the DREAM complex DNA binding protein LIN-54, the pocket protein that helps maintain the DREAM complex LIN-35, LIN-15B, or the H3K9me1/2 methyltransferase, MET-2, causes impaired chromatin compaction and ectopic germline gene expression in early development. This suggests that binding of LIN-15B, the DREAM complex, and methylation of H3K9 are each necessary for proper gene repression at germline gene loci in somatic tissue. Loss of any one of these proteins causes ectopic expression of germline genes and improper organismal development during temperature stress.

Chromatin is found in a generally unorganized and open state early in development (Politz et al., 2013; Yuzyuk et al., 2009). As development proceeds and differentiation occurs, chromatin becomes compacted and organized in a tissue specific manner (Mutlu et al., 2018; Politz et al., 2013; Yuzyuk et al., 2009). Organization of chromatin within the nucleus precedes upregulation of zygotic gene expression and is important for correct cell fate decisions (Gaertner et al., 2012; Politz et al., 2013; Yuzyuk et al., 2009). Poised Pol II accumulates at promoters in a stage specific, but not tissue specific manner (Gaertner et al., 2012). This means that when gene expression is upregulated at the maternal-to-zygotic switch, it occurs in a tissue-independent manner (Gaertner et al., 2012). Establishment of tissue specific repression, including establishment of repressive chromatin, must occur before global gene expression is upregulated in order to achieve proper cell fate (Gaertner et al., 2012; Poliltz et al., 2013; Yuzyuk et al., 2009). We hypothesize that repression by synMuv B may be a tissue specific mechanism to establish repressive chromatin environments before the maternal-to-zygotic switch. Loss of these proteins causes global changes in chromatin compaction into the developmental time of the maternal-to-zygotic switch, especially at high temperature, leaving their target loci vulnerable to expression when zygotic gene expression is upregulated.

Methylation of H3K9 is imperative to achieve chromatin compaction at the right time and place during development. Loss of MET-2 causes loss of ~80% of H3K9me1/2 across the genome. Previous studies have shown that *met-2* mutants ectopically express germline genes (Andersen and Horvitz, 2007, Towbin et al., 2012). *met-2* mutants also demonstrate the HTA phenotype (Rechtsteiner et al., 2018). Finally, it has also been shown that *met-2* mutants demonstrate decompacted chromatin in mid-to-late stages of embryogenesis (Fakhouri et al., 2010). In this study we show that *lin-15B, lin-54,* and *lin-35* mutants phenocopy *met-2* mutants in general chromatin compaction assays and at loci normally expressed in the germline in the intestinal lineage. Interestingly, the DREAM complex binds only ~1400 and LIN-15B binds to ~2100 locations throughout the genome. Loss of LIN-15B or DREAM binding, at relatively few places throughout the genome, causes the same phenotypes as losing ~80% of H3K9me1/2 in *met-2* mutants. One explanation for this similarity is that LIN-15B and the DREAM complex may be tissue specific regulators of H3K9 methylation at synMuv B regulated loci and this modification may be important for repression of germline genes in somatic tissue. Indeed, recent work has shown that LIN-15B is necessary for the specific localization of H3K9me2 to the promoters of germline genes that are misexpressed in synMuv B mutants (Rechtsteiner et al., 2019). This data, along with our work presented here, support a model in which LIN-15B promotes H3K9 methylation by MET-2 on germline genes at the correct stage of development for their repression in the soma. Although *lin-15B* mutants have exacerbated phenotypes at 26°C compared to 20°C, including decompacted chromatin, the loss of H3K9me2 on the promoters of germline genes in *lin-15B* mutants is not temperature sensitive (Rechtsteiner et al., 2018). Thus, loss of promoter H3K9me2 is not sufficient to explain the temperature sensitivity of synMuv B mutants, and the vulnerability of chromatin compaction to elevated temperatures requires further study.

Ectopic germline gene expression is increased in synMuv B mutants at 26°C compared to 20°C (Petrella et al., 2011). Before this study, the underlying mechanisms for this temperature sensitive phenotype have remained unclear. Here, we show that chromatin compaction is temperature sensitive, even in wild-type backgrounds, and may contribute to this phenotype. Chromatin has been shown to be reactive to changes in temperatures. Previous studies have reported that plants can change chromatin dynamics to modify gene expression in response to changes in temperature (Zografos and Sung, 2012). In addition to being temperature respondent, chromatin is also temperature sensitive. Early studies in *Drosophila* revealed that high temperature causes incomplete heterochromatin formation (Hartmann-Goldstein, 1967). This is the first study to investigate the role of temperature stress on chromatin compaction in a developmental context. Here we reveal that chromatin compaction in early development is especially vulnerable to the temperature sensitive nature of chromatin. Chromatin compaction is developmentally delayed when worms experience high temperatures for small amounts of time. Causing further stress to this system by loss of synMuv B proteins results in more drastic delays in chromatin compaction. Loss of these proteins not only cause changes in chromatin compaction at specific loci that they regulate, but also globally. Thus small changes in local chromatin domains throughout the genome, plus temperature stress, can indirectly change the chromatin landscape at a global level within the nucleus.

The experiments performed here were all done with a high temperature condition at the end of the temperature range where laboratory experiments are routinely done. This raises the question of whether these temperature stresses reflect levels that these organisms would experience under more natural conditions. Work in the last 10-15 years has greatly expanded our understanding of the ecology of naturally occurring *C. elegans* populations (Schulenburg and Felix, 2017). Unlike what was previously thought, it is now known that the reproductive lifespan of *C. elegans* occurs at the soil surface, often within rotting plant vegetation (Felix and Duveau, 2012; Frezal and Felix, 2015; Kiontke et al., 2011). In our temperature shift experiments, a window of 6-8 hours within embryogenesis at 26°C elicited a severe chromatin delay in developing embryos, even in wild-type. Given the short time period needed, it seems likely, that there are circumstances where *C. elegans* embryos in natural habitats would experience the temperatures tested here. Thus, having mechanisms that regulate chromatin compaction during development at these temperatures is necessary for species survival.

We found that delayed chromatin compaction in synMuv B mutants demonstrated an anterior-to-posterior pattern within intestinal cells. The anterior cells of the intestine were the last to adopt closed chromatin, a pattern independent of replication capacity, as there is a both final anterior and posterior division to form the *C. elegans* intestine. The *C. elegans* intestine is patterned anteriorly to posteriorly by Wnt signaling proteins (Fukushige *et al,* 1996; Lin *et al,* 1998; Schroeder and McGhee, 1998). Wnt signaling associated transcription factors are loaded into the most anterior cells of the intestine at each division (Schroeder and McGhee, 1998). It is possible that anteriorly loaded transcriptional factors are able to ectopically bind and express vulnerable uncompacted germline genes in the anterior intestine leading to high temperature arrest.

In this study, we demonstrate that synMuv B proteins are needed for chromatin compaction throughout development, especially at high temperature. Loss of chromatin compaction during developmental time periods in which germline genes are globally upregulated during the maternal-to-zygotic switch may underlie the developmental arrest phenotype of synMuv B mutants. It is important to understand how germline genes are repressed in somatic tissue as many somatic cancers that are highly proliferative and metastatic show upregulation of germline genes (Gure et al., 2005; Maine et al., 2016; Xu et al., 2014). Like *C. elegans,* mammals also repress germline gene expression during embryogenesis in somatic cells and recent work has indicated that this repression is dependent on H3K9me2 (Ebata, et. al., 2017; Lian, et. al., 2018). Given that the DREAM complex is completely conserved between worms and mammals (Litovchick, et al., 2007), its role in establishment of developmental repressive chromatin at germline genes may also be conserved. Further studies of synMuv B regulation will provide evidence of how organisms establish and maintain chromatin structure and gene expression states to ensure vitality.

## Materials and Methods

### Strains and nematode culture

*C. elegans* were cultured on NGM plates seeded with *E. coli* strain AMA1004 at 20°C unless otherwise noted. N2 (Bristol) was used as wild-type. Strain LNP0050 containing the transgene bnEx80(68xlacO+myo-3::mCherry+worm genomic DNA); gwls39[baf-1p::GFP::lacI::let-8583\’UTR;vit-5::GFP];caIs79(elt-2p::dTomato;pRF4)) was created for this study and crossed into mutant worms. Mutant alleles used were *lin-15B(n744), lin-35(n745), lin-54(n2231), lin-37(n758),* and *met-2(n4256).* The complete strain list can be found in supplemental table 1.

### Nuclear Spot Assay

Wildtype and mutant embryos and L1s with gwls39[baf-1p::GFP::lacI;;let-858 3\’UTR; vit-5::GFP]; bnEx85 (68xlacO+myo-3::mcherry+genomic C05C10.7+worm genomic DNA)and caIs79[elt-2p::dTomato, pRF4 (rol-6+)] arrays were collected and fixed with methanol and acetone and mounted using Gelutol (Petrella et al., 2011). Each genotype was scored at 8E, 16E, comma, and L1. High temperature samples were collected by upshifting L4 P0 worms to 26°C before collecting F1 embryos or L1s. L1 larvae were obtained by hatching embryos in the absence of food in M9 buffer. Images were acquired in Z-stacks using a Nikon Inverted Microscope Eclipse Ti-E Ti-E/B confocal microscope at 100X. For chromatin compaction analysis, each intestinal cell’s array was scored by eye as either open or closed as described in Yuzyuk *et al,* 2009. Nuclear volumetric array compaction analysis was performed using the isosurface function in Metamporph V7.8.8.0. Specifically the volume of both the nucleus and GFP+ LacO arrays of each individual intestinal cell was measured and used to determine the percentage of the nucleus that the array represented. Statistical analysis was performed by 2-way ANOVA using Prism Graph Pad.

### embryo E cell location lacO array scoring

Embryos scored for chromatin compaction analysis were also scored for percentage of open cells within an embryo. The intestinal cells at each developmental stage were lettered anterior-to-posterior as in WormBook (McGhee, 2007) so that chromatin compactions in each specific intestinal cell could be compared between embryos. The exact intestinal cells with open arrays in each embryo were recorded and the percentages of open intestinal cells were calculated. The percentage of each cell as open was calculated to determine the location of open cells within the embryo intestine. Statistical analysis was performed by 2-way ANOVA using Prism Graph Pad.

### Fluorescence In Situ Hybridization

Fluorescent probes were made by PCR labeling as described in Meister *et al.,* 2013 using fluorescent 568-5-dUPT (Thermo Fisher C11399) and ULYSIS Nucleic Acid Labeling Kits using Alexa Fluor 488 (molecular probes life technologies U21650). Worm genomic DNA was used as the template to create five ~500bp probes ~50kbp upstream and downstream of *pha-4* and *myo-3* as controls and *ekl-1, coh-3,* and C05C10.7 experimental loci. High temperature samples were collected by upshifting L4 P0 worms to 26°C before collecting F1 embryos. Embryos were dissected in 1X M9 on slides and then frozen in liquid nitrogen. After freeze-cracking, slides were fixed in methanol (on ice, 2 minutes) followed by 4% formaldehyde (4°C, 10 minutes) and then washed two times in 1X PBS (room temperature, 2 minutes). Embryos were permeabilized in 0.5% PBS-Triton X-100 (room temperature, 5 minutes), washed two times in 1XPBS (room temperature, 2 minutes), rinsed in 0.01 N HCl, and incubated in 0.1 N HCl (room temperature, 2 minutes). Slides were then washed once with 1X PBS (room temperature, 3 minutes), once with 2X SSC (room temperature, 3 minutes) and then treated with 50 ug/mL of RNase A (VWR 1247C393) in 2X SSC for 45 minutes at 37°C by overlaying the sample with 500ul of RNAse solution and incubating in a humidified chamber. Slides were then washed once with 2X SSC (room temperature, 2 minutes) and than incubated in 2X SSC/50% formamide for 2 hours at room temperature. The probe containing sample was diluted to a concentration of 2-5 ng/ul in 100% deionized formamide and an equal volume of hybridization buffer was added. 25 ul of the probe solution was added to the sample and overlayed with a glass chamber sealed with cement glue. The slides were pre-hybridized overnight at 37°C in a humidified chamber. FISH probe and sample were denatured at 76°C by placing the slides on a heat block and then incubated for hybridization for 3 days at 37°C. Hybridization was followed by three washes of 2X SSC (37°C, 5 minutes) and two times in 0.2X SSC (55°C, 5 minutes). Images were captured on Nikon Confocal and analyzed using Metamorph isosurface function to determine the 3-D distance between the two centroids as described in Yuzyuk et al., 2009. Statistical analysis was performed by 2-way ANOVA using Prism Graph Pad.

### Embryo Shift Assays

For downshifting experiments: P0 L4 wildtype and mutant strains expressing *caIs79[elt-2p::dTomato, pRF4 (rol-6+)]* worms were upshifted to 26°C. The next day, 2-cell embryos were dissected out of adults in 1X M9, plated on NGM, and the plate was returned to 26°C. Developmental stages were determined based on the number of intestinal cells made visible by the *caIs79* transgene. Embryos were downshifted to 20°C when the majority of embryos on the plate were at 4E (~2 hours), 8E (~3 hours), 16E (~5 hours), comma (~7 hours), pretzel ~(9 hours), and L1 (~14 hours). The HTA phenotype was scored two days after shifting as described in Petrella et al., 2011. Each downshift was done 4-6 times with 10-20 embryos plated per replicate. For upshifting experiments a similar protocol was used: 2-cell embryos were dissected out of adults and maintained continually at 20°C. After dissection embryos were plated and kept at 20°C until being upshifted to 26°C at 4E, 8E, 16E, comma, pretzel, and L1 stage. Plates were scored for HTA two days after shifting.

### PGL expression analysis

P0 L4 wildtype and mutant strains expressing *caIs79[elt-2p::dTomato, pRF4 (rol-6+)]* and *pgl-1(sam33[pgl-1::gfp::3xFlag])* worms were upshifted to 26°C. The next day, adults were dissected in 1X M9 and 2-cell embryos placed in 1X M9 on polylysine coated slides in humid chambers and returned to 26°C. Embryos were downshifted on slides to 20°C at 8E, 16E, and comma stage. Arrested L1 larva were fixed in methanol and acetone and imaged in Z stack using a Nikon Inverted Microscope Eclipse Ti-E Ti-E/B confocal microscope at 60X. Mean fluorescent intensity of PGL-1:GFP was determined using ImageJ. Statistical analysis was performed by 2-way ANOVA using Prism Graph Pad.

## Acknowledgments

Thanks to Jim McGhee for JM163 strain and Dustin Updike for DUP75. Some strains were provided by the CGC, which is funded by NIH Office of Research Infrastructure Programs (P40 OD010440). Many thanks to Anita Manogaran for comments and discussion of the manuscript. This work was supported by a NIH grants R00GM98436 and R15GM122005 to L.N.P.

