## Supplemental Figures and Table for "*C. elegans* synMuv B proteins regulate spatial and temporal chromatin compaction during development"

Figure S1

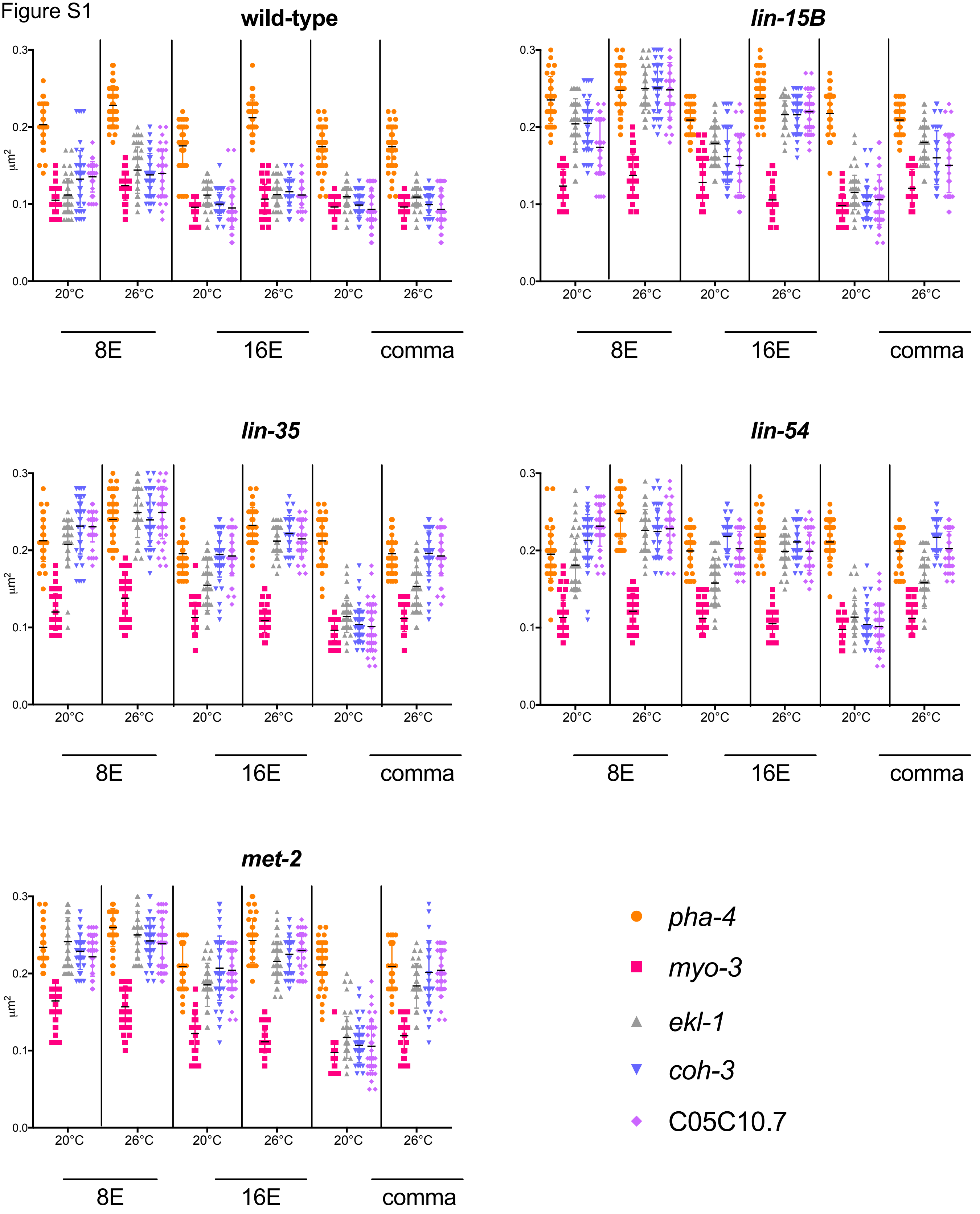
wild-type
lin-15B
lin-35
lin-54
met-2
μm2
0.0
0.1
0.2
0.3
20°C
26°C
8E
16E
comma
pha-4
myo-3
ekl-1
coh-3
C05C10.7

Figure S2

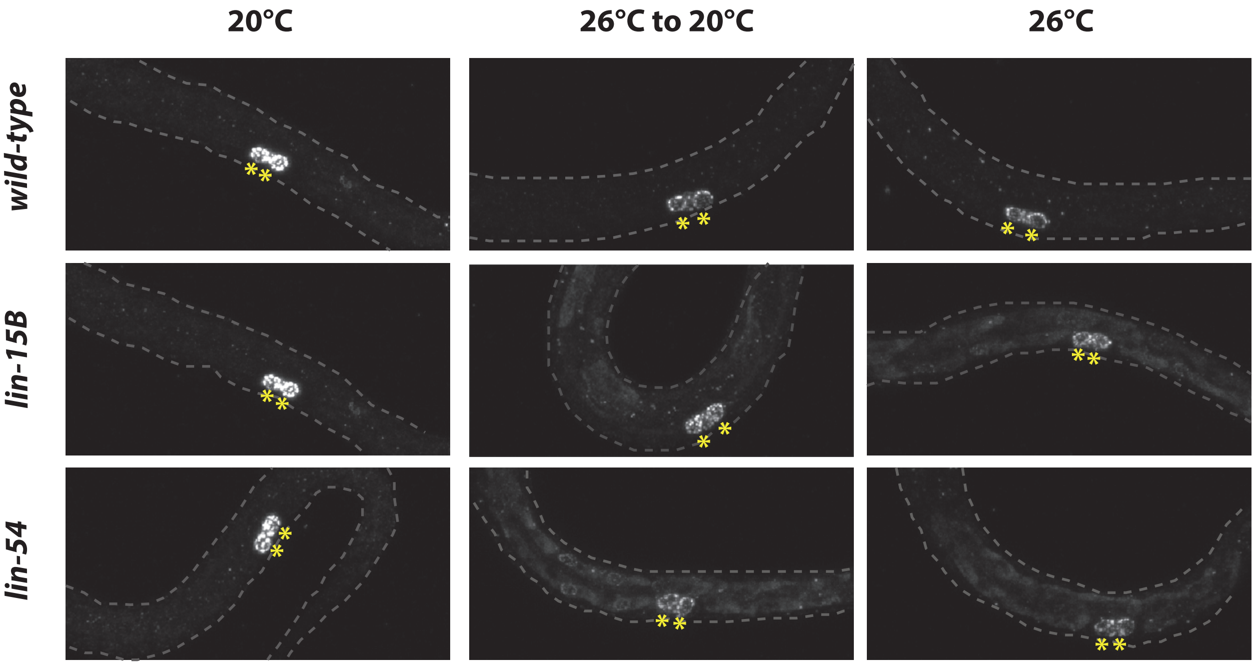

**Figure S1 Related to Figure 4: Fluorescent *in-situ* hybridization of synMuv B regulated loci reveals more open chromatin in synMuv B mutants at higher temperatures**

Wild-type (A), *lin-15B* (B), *lin-35* (D), *lin-54* (D), and *met-2* (E) embryos at 20°C and 26°C at 8E, 16E and comma stage were labeled with 568-5-dUTP (red) and Alexa Fluor 488 (green) probes 50kb upstream and 50kb downstream of *myo-3, pha-4, ekl-1, coh-3* and C05C10.7. Each dot represents the 3-D-distances between centroids ($\mu m^2$) of one intestinal cell.

**Figure S2 Related to Figure 5: synMuv B mutant embryos downshifted to low temperature at 16E ectopically express PGL-1**

Wild-type, *lin-15B* mutant, and *lin-54* mutant embryos carrying a PGL-1:GFP transgene were placed at 26°C or 20°C in 1XM9 drops. Embryos were either kept at 26°C, downshifted to 20°C at 16E, or maintained at 20°C, and allowed to arrest at the L1 stage. L1 worms were fixed and imaged in Z-stack using confocal microscopy. Panels represent maximum projection of PGL-1:GFP. Asterisks mark primordial germ cells.

**Table S1**

| Strain Name | Genotype | Source |
|---|---|---|
| DUP0075 | *pgl-1(sam33[pgl-1::gfp::3xFlag])* | Dustin Updike |
| JM163 | caIs79[elt-2p::dTomato, pRF4 (rol-6+)] | Jim McGhee |
| LNP0024 | *lin-35(n745);* bnEx80(68XlacO +myo-3::mCherry+ worm genomic DNA); gwIs39[baf-1p::GFP::lacI::let-858 3\'UTR; vit-5::GFP] | This Study |
| LNP0026 | *lin-54(n2231);* bnEx80(68XlacO +myo-3::mCherry+ worm genomic DNA); gwIs39[baf-1p::GFP::lacI::let-858 3\'UTR; vit-5::GFP] | This Study |
| LNP0027 | *met-2(n4256);* bnEx80(68XlacO +myo-3::mCherry+ worm genomic DNA); gwIs39[baf-1p::GFP::lacI::let-858 3\'UTR; vit-5::GFP] | This Study |
| LNP0038 | caIs79(elt-2p::dTomato;pRF4); *pgl-1(sam33[pgl-1::gfp::3xFlag])* | This Study |
| LNP0040 | *lin-15B(n744)* bnEx80(68XlacO +myo-3::mCherry+ worm genomic DNA); gwIs39[baf-1p::GFP::lacI::let-858 3\'UTR; vit-5::GFP]; (caIs79(elt-2p::dTomato;pRF4)) | This Study |
| LNP0041 | *lin-15B(n744)* caIs79(elt-2p::dTomato;pRF4); *pgl-1(sam33[pgl-1::gfp::3xFlag])* | This Study |
| LNP0049 | *lin-54(n2231)* caIs79(elt-2p::dTomato;pRF4); *pgl-1(sam33[pgl-1::gfp::3xFlag])* | This Study |
| LNP0050 | bnEx80(68xlacO+myo-3::mCherry+worm genomic DNA); gwIs39[baf-1p::GFP::lacI::let-8583\'UTR;vit-5::GFP];caIs79(elt-2p::dTomato;pRF4)) | This Study |
